# 10-Month-Old Infants Are Sensitive to the Time Course of Perceived Actions: Evidence From a Study Combining Eye-tracking and EEG

**DOI:** 10.1101/131516

**Authors:** Cathleen Bache, Anne Springer, Hannes Noack, Waltraud Stadler, Franziska Kopp, Ulman Lindenberger, Markus Werkle-Bergner

## Abstract

Research has shown that infants are able to track a moving target efficiently – even if it is transiently occluded from sight. This basic ability allows prediction of when and where events happen in everyday life. Yet, it is unclear whether, and how, infants internally represent the *time course* of ongoing movements to derive predictions. In this study, 10-month-old crawlers observed the video of a same-aged crawling baby that was transiently occluded and reappeared in either a temporally *continuous* or non-continuous manner (i.e., *delayed* by 500 ms vs. *forwarded* by 500 ms relative to the real-time movement). Eye movement and rhythmic neural brain activity (EEG) were measured simultaneously. Eye movement analyses showed that infants were sensitive to slight temporal shifts in movement continuation after occlusion. Furthermore, brain activity related to sensorimotor rather than mnemonic processing differed between observation of continuous and non-continuous movements. Early sensitivity to an action’s timing may hence be explained within the internal real-time simulation account of action observation. Overall, the results support the hypothesis that 10-month-old infants are well prepared for internal representation of the time course of observed movements that are within the infants’ current motor repertoire.

## 1. Introduction

Infants possess a remarkable ability to predict future events. This has been demonstrated in various domains such as visual expectation (Canfield and Haith, 1991;Adler et al., 2008), social interaction (Adamson and Frick, 2003;Striano et al., 2006), action perception (Hunnius and Bekkering, 2010;Rosander and von Hofsten, 2011), and object tracking (Rosander and von Hofsten, 2004). Predicting when and where events occur is indispensable to understand and smoothly coordinate one’s behavior with others’ actions in everyday life (cf. Hommel et al., 2001). However, it is unclear whether infants actually rely on real-time processing of observed actions when predicting their future trajectory. As a consequence, the cognitive and neural processes of such real-time representations remain poorly understood.

Transient occlusion of ongoing movement is a frequently used paradigm to investigate predictive abilities and their neural implementations. According to this research, both *mnemonic* processes (Wilcox and Schweinle, 2003;Keane and Pylyshyn, 2006;Bosco et al., 2012;Springer et al., 2013) and *sensorimotor* processes (e.g., Graf et al., 2007;Southgate et al., 2009;Elsner et al., 2013) have been advocated to assist movement observation. Studies on *object motion* suggest that infants linearly extrapolate the ongoing trajectory of observed movement (e.g., von Hofsten et al., 1998). Linear extrapolation corresponds to working *memory* operations (e.g., Baddeley and Hitch, 1974;Pelphrey and Reznick, 2002) maintaining an internal representation of the target movement during occlusion that can be matched following the reappearance to generate predictions. In line with this assumption, infants need to plan and control their eye movements based on previously collected information in order to match pre-and post-occlusion input (Bennett and Barnes, 2003;Rosander and von Hofsten, 2004;Springer et al., 2013;Kwon et al., 2014;Bache et al., 2015).

While object motion usually follows linear trajectories with continuous velocity human movement is non-linear with changes in velocity and path. Linear extrapolation may hence not be an optimal approximation of human trajectories. Infants have been shown to render precise predictions about observed *human actions*, such as transporting a ball into a basket. Here, predictions may be derived from *internally simulating* the observed action in sensorimotor areas of the brain as if performing the action oneself (Flanagan and Johansson, 2003;Falck-Ytter et al., 2006;Rosander and von Hofsten, 2011). In line with this assumption, initial evidence suggests that *sensorimotor* processes support the internal representation of spatiotemporal aspects of human action in infants, including predictive functions (Southgate et al., 2009;Southgate et al., 2010;Stapel et al., 2010;Stapel et al., 2016).

It remains unclear whether infants’ processing of human movement recruits *real-time* representations employing simulation, memory, or both. Here, we consider representations as a neural pattern of stimulus coding that maintains stimulus properties as a close analogue to the original sensory input in order to integrate previous and newly incoming stimulation (Hebb, 1949/2009).

Transient occlusion allows manipulating the temporal structure of on-going movement so that the post-occlusion trajectory does not reflect a time-matching continuation of the pre-occlusion movement. Applying such a paradigm, behavioral studies in adults pointed out that the processing of observed actions is running parallel to the actions’ time course (e.g., Graf et al., 2007). However, previous studies also suggested that delayed and forwarded manipulations may not be processed similarly. More precisely, adults judged the continuation of a human action after a transient occlusion to be continuous when it was in fact slightly delayed, while they judged the continuation to be on time when it was in fact slightly forwarded (e.g., Sparenberg et al., 2012). Infants could recognize temporal shifts only if extreme jumps forward in time were presented (Wilcox and Schweinle, 2003;Bremner et al., 2005), while they could readily detect an one-second delay in their mothers’ interaction (Striano et al., 2006). To further explore how infants process the time course of human action, delayed *and* forwarded movements need to be contrasted with continuous movement.

The present study aimed to investigate infants’ sensitivity to the time course of human action. Specifically, 10-month-old crawlers watched a same-aged crawling baby that was transiently covered from sight. Following the occlusion, the movement was either continued in a time-matching manner (i.e., no time shift, resulting in continuous movement continuation) or in a non-matching manner (i.e., time shift, resulting in delayed or forwarded movement continuation) relative to the pre-occlusion movement stream (Graf et al., 2007). Due to limits in attention span, infants were randomly assigned to one of two experimental groups watching either *continuous* and *delayed* (i.e., Delay group) or *continuous* and *forwarded* movements (i.e., Forward group) within a single experimental session.

To capture mnemonic and sensorimotor contributions to movement processing, eye movements (via eye-tracking) and rhythmic neural activity (via electroencephalography, EEG) were measured simultaneously. Eye movements have been associated with both mnemonic (e.g., Keane and Pylyshyn, 2006) and sensorimotor processing (e.g., Elsner et al., 2013) and therefore provide a rather indirect measure of cognitive processes. Rhythmic neural activity may provide a complementary view. Specifically, *mnemonic functions* are assumed to be reflected in *frontal theta* modulations (Jacobs and Kahana, 2010;Saby and Marshall, 2012;Lisman and Jensen, 2013;Bache et al., 2015), and *sensorimotor simulation* is assumed to be reflected in *central alpha* modulations (also labeled sensorimotor, rolandic or mu rhythm; Cochin et al., 1999;Muthukumaraswamy et al., 2004;Marshall et al., 2011;Bache et al., 2015).

Only if the ongoing movement was processed in real-time while it was hidden during occlusion, could a time-matching continuation be distinguished from a non-matching one following occlusion (cf. Graf et al., 2007). Hence, infants’ sensitivity to the time course of movements would be reflected in differences in tracking and neural patterns following occlusion, whereas there should be no differences prior to and during the occlusion. With regard to *eye-tracking*, we hypothesized that the tracking of the target’s reappearance position would be more accurate (i.e., landing on mid to front parts of the target) and more consistent (i.e., less variable across infants) in time-matching continuations. In contrast, the reappearance position would be overshot (i.e., landing in front of the target) in delayed continuations, and undershot (i.e., landing behind the target) in forwarded continuations, and tracking would be overall less consistent in both non-continuous continuations. With regard to *EEG*, we hypothesized that *frontal theta* activity would be elevated more when processing non-matching than when processing time-matching continuations because temporarily stored representations during occlusion would not match the reappearance position following occlusion (Orekhova et al., 1999;Kwon et al., 2014). Secondly, *central alpha* activity was expected to decrease more in non-matching than in time-matching continuations because real-time simulation during occlusion should result in a prediction error relative to the actual reappearance position following occlusion (Kilner et al., 2007;Stapel et al., 2010).

## 2 Methods

### 2.1 Participants

Participants were recruited from a database of parents interested in participating in infant studies at the Max Planck Institute for Human Development, Berlin. Infants were invited at 10 months of age (± 10 days) according to the following criteria: *(a)* the infant was born at term (week of gestation ≥ 37, birth weight ≥ 2500 g), *(b)* to the parents’ knowledge, the infant had no visual impairments nor current health issues, and *(c)* according to the parents, the infant was capable of crawling on hands and knees with her/his stomach lifted but not yet able to walk. Parents were encouraged to bring their own notes about their children’s motor development to fill in a short checklist in the lab. The experiment was approved by the Institute’s Ethics Committee.

A total of 99 10-month-old infants were tested. Twelve infants were not considered for further preprocessing as they did not crawl a distance of 1.5 m in the lab at least once (n = 4) or were too fussy to be properly tested following preparation for EEG and eye-tracking (n = 8). For *eye-tracking analysis*, 14 further infants were excluded because *(a)* the calibration failed (n = 3), *(b)* the trigger information was missing in the recorded data (n = 6), *(c)* the measurement failed due to technical issues (n = 4), or *(d)* fewer than 10% of the actually watched trials were free of artifacts (n = 1). Furthermore, for the *EEG analysis*, 37 further infants were excluded because they did not produce enough artifact-free EEG data (at least 10 trials per condition; n = 30) or the measurement failed due to technical issues (n = 7).

Thus, the final eye-tracking sample consisted of **32** infants in the Delay group and **31** infants in the Forward group, and the final EEG sample comprised **24** infants in the Delay group and **25** infants in the Forward group. *Table 1* and *Table 2* provide descriptive information on the final samples for eye-tracking and EEG analysis, respectively. Figure 1 illustrates which trials of both eye-tracking and EEG data were contributed to the analysis within the final samples. Note that not all infants provided data in both measures, and artifact-free trials were contributed randomly throughout the test session. As a result, eye-tracking and EEG data were analyzed separately (cf. Stapel et al., 2010).

**Table 1.** Descriptive information on eye-tracking sample.

|  | Delay group (*N* = 32) |  | Forward group (*N* = 31) |  |
| --- | --- | --- | --- | --- |
|  | *M* | *SD* | *M* | *SD* |
| Number of boys/girls | 20/12 |  | 15/16 |  |
| Age in days<sup>1</sup> | 298.2 | 5.8 | 301.2 | 6.1 |
| Week of gestation at delivery | 39.8 | 1.3 | 40.3 | 1.0 |
| Birth weight in grams | 3385 | 390 | 3606 | 373 |
| Onset age in months<sup>2</sup> |  |  |  |  |
| crawling | 8.0 | 1.1 | 7.9 | 0.9 |
| sitting | 7.6 | 1.2 | 7.1 | 1.5 |
| standing | 8.5 | 0.9 | 8.2 | 0.9 |
| Number of trials<sup>3</sup> |  |  |  |  |
| continuous movement | 13 | 8.8 | 11 | 7.5 |
| time-shifted movement<sup>4</sup> | 11 | 8.2 | 11 | 8.3 |

*Note.* *M* = mean, *SD* = standard deviation. Participants were randomly assigned to one of the two experimental groups (i.e., Delay group vs. Forward group). <sup>1</sup> 300 days equals 10-month birthday. <sup>2</sup> According to parents' report. <sup>3</sup> Available for analysis after preprocessing. <sup>4</sup> Delayed in the Delay group and forwarded in the Forward group. In the Delay group, one child had not yet mastered sitting independently, another pulling up in a standing position; in the Forward group, one child had not yet mastered sitting independently and two children were not yet able to pull themselves up in a standing position. Exceptions were not the same children and thus not excluded. Mean age ( $t = -2.2, p = .033$ ) and birth weight was lower in the Delay group ( $t = -2.3, p = .025$ ); no further differences were found ( $t < -1.9, p > .051$ ). EEG and eye-tracking samples (see also *Table 2*) did not differ ( $t < 1.5, p > .150$ ) except for the number of trials ( $t = 3.4, p = .001$ ) due to varying inclusion criteria for eye-tracking and EEG data.

**Table 2.** Descriptive information on EEG sample.

|  | Delay group (*N* = 24) |  | Forward group (*N* = 25) |  |
| --- | --- | --- | --- | --- |
|  | *M* | *SD* | *M* | *SD* |
| Number of boys/girls | 9/15 |  | 12/13 |  |
| Age in days | 296.8 | 5.6 | 301.0 | 5.8 |
| Week of gestation at delivery | 39.5 | 1.5 | 40.0 | 1.0 |
| Birth weight in grams | 3327 | 373 | 3569 | 318 |
| Onset age in months |  |  |  |  |
| crawling | 8.0 | 1.1 | 8.1 | 0.7 |
| sitting | 7.6 | 1.1 | 6.9 | 1.6 |
| standing | 8.5 | 0.7 | 8.5 | 0.7 |
| Number of trials |  |  |  |  |
| continuous movement | 21 | 7 | 19 | 8 |
| time-shifted movement | 20 | 9 | 21 | 8 |

*Note.* Information is shown in analogy to Table 1. In the Delay group, two children had not yet mastered sitting independently, another child pulling up in a standing position; in the Forward group, one child had not yet mastered sitting independently and two children were not yet able to pull themselves up in a standing position. Exceptions were not the same children and thus not excluded. Age ( $t = -2.6, p = .014$ ) and birth weight ( $t < -2.5, p = .018$ ) were lower in the Delay group; no further differences were found ( $t < -1.8, p > .075$ ).

**Figure 1.**
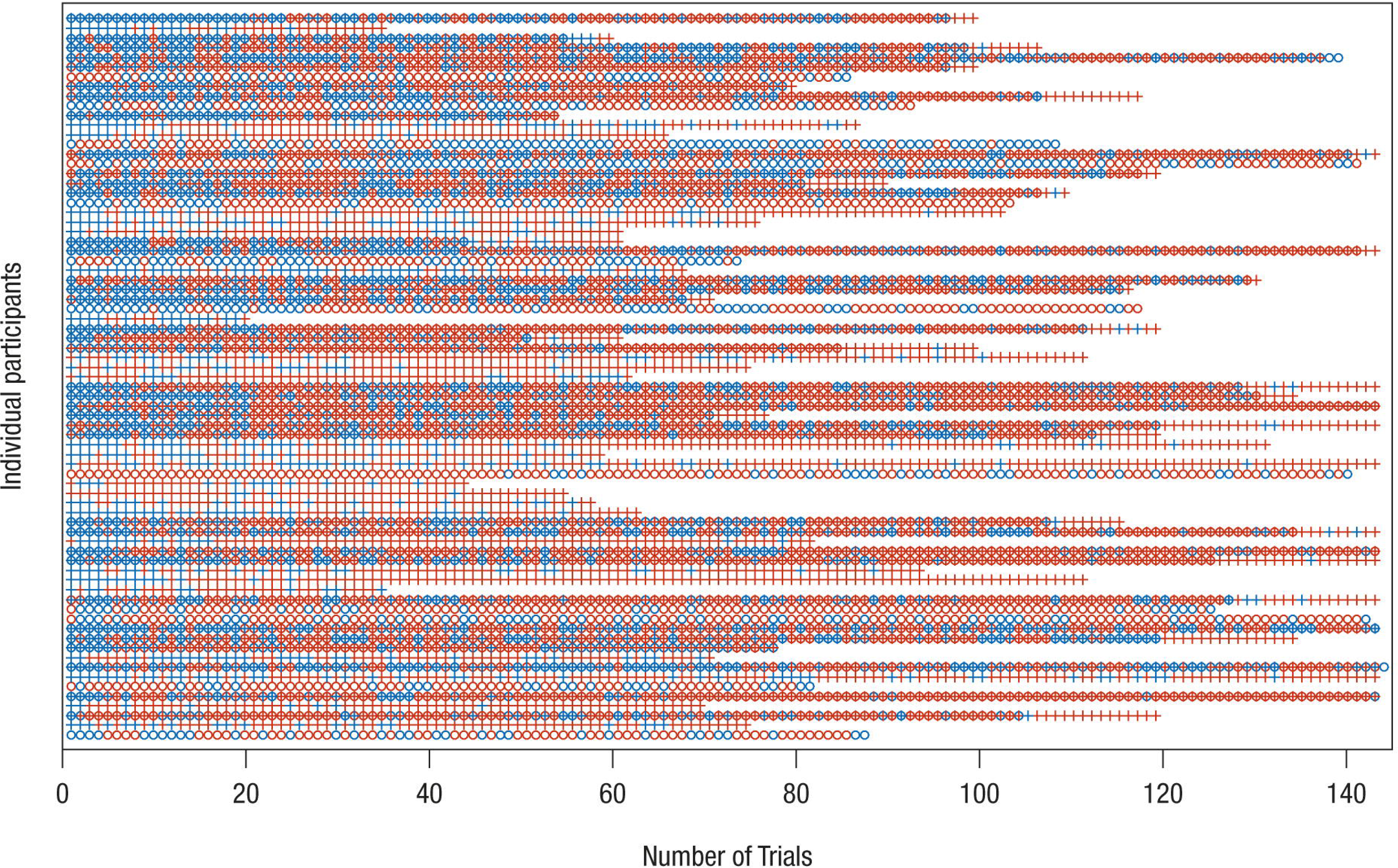
Distribution of trials included in analysis of EEG and eye-tracking data. On the y-axis, each row represents one data set/participant; only participants who were included in the final sample are shown. The x-axis shows the chronological trial number. Blue – trial available for analysis; red – trial not available for analysis. Circle – EEG data, Cross – eye-tracking data. Note that not for all data sets measurement of both EEG and eye-tracking was possible. It is apparent that infants contributed trials to the final analysis more or less randomly. Therefore, separate analyses of eye-tracking and EEG measures were performed.

### 2.2 Stimulus material and procedure

Participants repeatedly watched a video of a same-aged baby crawling in front of a light gray background (2480 ms; *pre-occlusion phase*). The baby’s movement was transiently occluded by a full-screen black occlusion (500 ms; *occlusion phase*) and then immediately continued (1000 ms; *post-occlusion phase*). Hence, each trial lasted for 4000 ms. The video however was 4500 ms long, allowing to manipulate the movements’ timing. We choose to present an intransitive movement, that is a movement not directed at an apparent object or goal, in order to avoid confounds with object knowledge or object saliency. To avoid lateralization of brain activity, each video was presented from both left to right and right to left (i.e., flipped versions of the original video). On the x-axis of the monitor, the stimulus (i.e., crawling baby) was on average 279 pixel (ranging from 207 to 315 pixel) wide and moved with an average speed of 3° visual angle per second (see Figure 2 for an illustration of the stimulus material).

**Figure 2.**
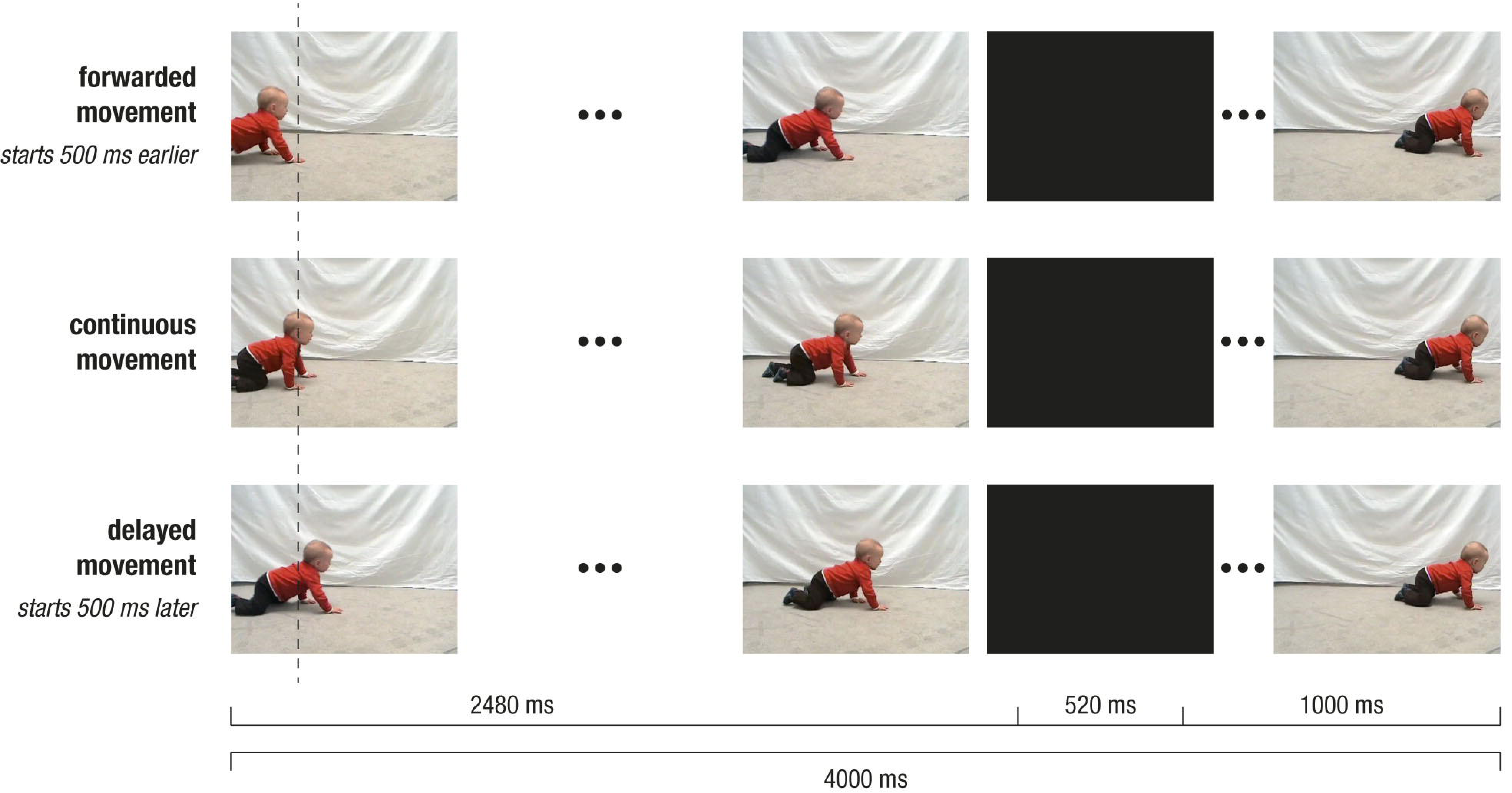
Depiction of stimulus design. Screenshots of crawling movement at pre-occlusion, occlusion, and post-occlusion phases, for continuous movement (middle row), forwarded movement (upper row) and delayed movement (lower row). Note that, during pre-occlusion, the starting time in the video clip depended on the experimental condition: The continuous movement started at 500 ms, the delayed movement at 1000 ms and forwarded movement at 0 ms. Therefore, movement positions slightly differed across conditions as indicated by the vertical dotted line. Following occlusion, the video was always continued with the same frame in the video (i.e., at 3000 ms), and therefore the visual input was identical across conditions.

In a between-subjects design, participants were randomly assigned to one of two experimental groups: In the Delay group, *continuous* and *delayed* movements were shown, while in the Forward group, *continuous* and *forwarded* movements were presented. To achieve continuous and non-continuous (i.e., delayed or forwarded) movements, the starting time in the video footage was varied. More precisely, during pre-occlusion, non-continuous trials started either 500 ms earlier (i.e., at 0 ms in forwarded conditions) or 500 ms later (i.e., at 1000 ms in delayed conditions) as compared to the continuous trials (i.e., at 500 ms). However, following the occlusion (i.e., 500 ms), the movement was always continued at 3000 ms in the video footage. In other words, during occlusion, the video footage was paused in delayed trials (i.e., 0 ms elapsed), fast-forwarded in forwarded trials (i.e., 1000 ms elapsed), and continued in real-time in continuous trials (500 ms elapsed). Therefore, in non-continuous trials, the post-occlusion movement did not match a natural continuation of the pre-occlusion movement, but resulted in a forwarded (i.e., 500 ms too early) or a delayed (i.e., 500 ms too late) time course of the movement. Notably, the visual input slightly varied during pre-occlusion phases, while it was identical during occlusion and post-occlusion phases. Within each trial, time manipulation could only be detected following occlusion. This design ensured that differences between conditions during occlusion and post-occlusion could not be attributed to visual differences but reflect the manipulation of the movements’ time course.

Stimuli were presented using a customized program written in Microsoft Visual C++ (Microsoft Corporation, Redmond, USA). Each trial was preceded by a centered fixation object (i.e., colored pictures of toys; duration of 800 – 1300 ms) on gray background. Conditions were presented in blocks of six trials, because rapid learning over trials has been reported (see Henrichs et al., 2014). The order of blocks was quasi-randomized such that blocks with the same condition and movement direction were never repeated successively. Participants were randomly assigned to one of two predefined block orders per experimental group. The stimulus presentation was controlled by an experimenter; depending on infants’ attention and compliance up to 24 blocks (i.e., 144 trials) were presented. The experiment was conducted in an acoustically and electromagnetically shielded room. Experimental sessions were video-recorded in time-synchronized split-screen images including a frontal and lateral view of the infant as well as a running and a condition trigger for coding infants’ behavior post-hoc (Interact; Mangold International GmbH, Arnstorf, Germany). The lighting conditions were kept comparable across participants. The infant sat on the parent’s lap in a BabyBjörn® baby carrier facing a 20.1’’ monitor (dimensions: 40.8 cm × 30.6 cm, visual angle ≈ 29° × 22°) at a distance of approximately 80 cm (for more detailed information on the experimental procedure, see Bache et al., 2015). Despite restricting infant’s position, sitting distance could range from 60 cm to 90 cm when infants leaned forward or backward. In our set-up, the size of one pixel (0.051 cm) equals 0.037° visual angle for an ideal sitting distance.

### 2.3 Data acquisition

#### 2.3.1 Eye-tracking data

##### 2.3.1.1 Recording

Eye movements were recorded continuously using an EyeLink 1000 remote system eye-tracker (SR Research, Ottawa, Canada), which allows for free head movements. The eye-tracking camera including the infra-red source was permanently positioned centrally below the presentation monitor. Participants were seated 55 cm from the recording eye-tracking camera. The camera recorded the corneal relative to the pupil reflection of the left eye at a frequency of 250 Hz in terms of raw gaze positions in pixel.

The infants’ head position was tracked using a small sticker on their forehead that allowed accounting for head movement of up to 100 cm/s. Infants’ position relative to the head box of the eye-tracker was checked using the camera image before the experimental procedure started. The data were filtered online using the second stage of the built-in heuristic filter (Stampe, 1993) which reduces noise in the data by a factor of 4 to 6 (according to the EyeLink manual). The average accuracy of the eye-tracking system is 0.5° visual angle for an ideal participant (i.e., sitting still with minimal head movements and generating a perfect calibration), as reported by the providing company, which would approximate to a 0.07 cm area at the viewing distance of 80 cm in the present experiments.

Following EEG preparation and prior to stimulus presentation, a five-point calibration procedure on a gray background was performed in the following order: center, upper center, lower center, left center, right center. The calibration target was a dancing rabbit in a square shape (96 × 96 pixel, approximately 4.9 cm^2^ on the monitor and 3.5° visual angle from the sitting position) accompanied by an attractive sound. An experimenter pushed a button to accept the gaze position if it was on the target position. The central position was repeated at the end as an estimate of accuracy. Calibration was only accepted if it was reported to be ‘good’ by the recording software (i.e., average error < 1° visual angle) and if the overall pattern of gaze positions matched the target’s positions according to the experimenter’s evaluation. If the calibration was not accepted, it was repeated until it was satisfying. If calibration could not be obtained, the experimental procedure was continued, but the participants’ eye-tracking data were discarded from analysis.

##### 2.3.1.2 Preprocessing

Ideal preprocessing of eye-tracking data should yield data that represent artefact-free and task-relevant eye movement. Yet, in infant studies, raw eye-movement data are typically only preprocessed in terms of detecting saccades or fixations by applying built-in algorithms of the eye-tracking system at hand (e.g., Gredebäck and Melinder, 2010). Recently, Wass et al. (2014) demonstrated that data quality affects fixation detection to such an extent that the interpretation of the results is put into question – even when a satisfactory calibration outcome is achieved. Moreover, comparing common categorization algorithms, it has been shown that results for fixations and saccades vary to such an extent that automated categorization may not always return meaningful results (Komogortsev et al., 2010;see Wass et al., 2013, for calculation of data quality post-hoc).

In order to avoid classification artifacts and to account for data quality, raw gaze positions (i.e., x-and y-value in pixel per measurement unit) were visually inspected using a custom-made graphical user interface (GUI, see Supplementary Material) in MATLAB 7.10.0 (MathWorks Inc., Natick, MA, USA) to detect trials with *measurement errors* (i.e., noisy or no data, e.g., following gross movement, substantial changes in body/head position, or changes in the eyes’ lubrication) and *compliance failure* (e.g., gazing away from or staring blankly at the monitor; see Haith, 2004;Schneider et al., 2008;Wass et al., 2014). More precisely, raw data were segmented into 3400 ms long epochs from -2200 ms to 1200 ms relative to the onset of occlusion. The first and last 300 ms of each trial were discarded from analysis because *(a)* following stimulus onset, infants reoriented from the centered fixation object to the stimulus movement starting on either the left or right side of the monitor, and *(b)* approaching stimulus offset, infants’ attention frequently terminated. The extracted segments were displayed neutral with respect to condition, movement direction, and test session to avoid confounding influence. The stimulus dimensions (i.e., x-and y-values in pixel) for each video-frame were derived using OpenCV (http://opencv.org/) by defining the color contrast separating colored stimulus and grayish background. Stimulus dimensions were included in the GUI to map gaze positions to actual stimulus position. Only trials with less than 50% missing data (incl. data points beyond the monitor) were considered for inspection.

Each trial was visually scanned by a trained rater (CB) according to the persistent or repeated presence of the following exclusion criteria: *(a)* missing gaze positions, gaze positions outside and/or on the borders of the monitor shortly before, during, and/or following the occlusion in order to make sure that transitions were actually perceived, *(b)* noisy and/or broken data resulting from technical error, *(c)* prolonged stationary data points reflecting blank stares without following of the stimulus movement. In principle, trials could be associated with more than one criterion. Missing or outlying data points at the beginning and end of the trial were not regarded as an exclusion criterion. Trials that were identified as being of poor quality were discarded from further analyses (see Supplementary Material). In ambiguous trials, video-recordings of the experimental session were used to inform the decision.

Following visual inspection, the percentage of trials available for eye-tracking analysis was calculated relative to the number of trials that the infant had actually watched during stimulus presentation, based on behavioral coding of video-recordings. Only data from infants providing at least 10% artifact-free trials were considered for further analyses.

##### 2.3.1.3 Analysis of gaze positions over time

As the movement was mainly evolving on the horizontal axis across time, only raw gaze positions (in pixel) on the x-dimension (Gx) were used. Within subjects, gaze positions were averaged per condition for each measurement point (i.e., every 4 ms). Data for movement from right to left were flipped, so all trials were available in the left-to-right direction. Data on either the y-and/or x-axis that were outside of the monitor’s dimensions were considered missing, and this was also applied to the corresponding gaze position on the other axis. Missing values were discarded before averaging.

The analysis focused on infants’ gaze behavior in reaction to the moving stimulus. However, it is difficult to quantitatively determine gaze relative to moving objects based on raw gaze positions. To relate gaze and stimulus position, the midpoint of the minimal and maximal x-value of the stimulus dimension per video frame (see 2.3.1.2) was determined as mean stimulus position (in pixel). Due to the biological characteristics of crawling (i.e., stretching and flexing of extremities), the stimulus dimensions vary from frame to frame and thus the mean stimulus position over time does not represent a linear movement (see black dotted line in Figure 3A). Following, at each measurement point, the respective mean stimulus position was subtracted from the raw gaze position, resulting in a difference score that reflects the *distance* between gaze position and stimulus position. Thus, if infants were looking at the front parts of the stimulus target (i.e., baby’s hands and head), the resulting scores would be positive (and vice versa). Resulting difference scores were averaged for each measurement point per condition within each participant.

**Figure 3.**
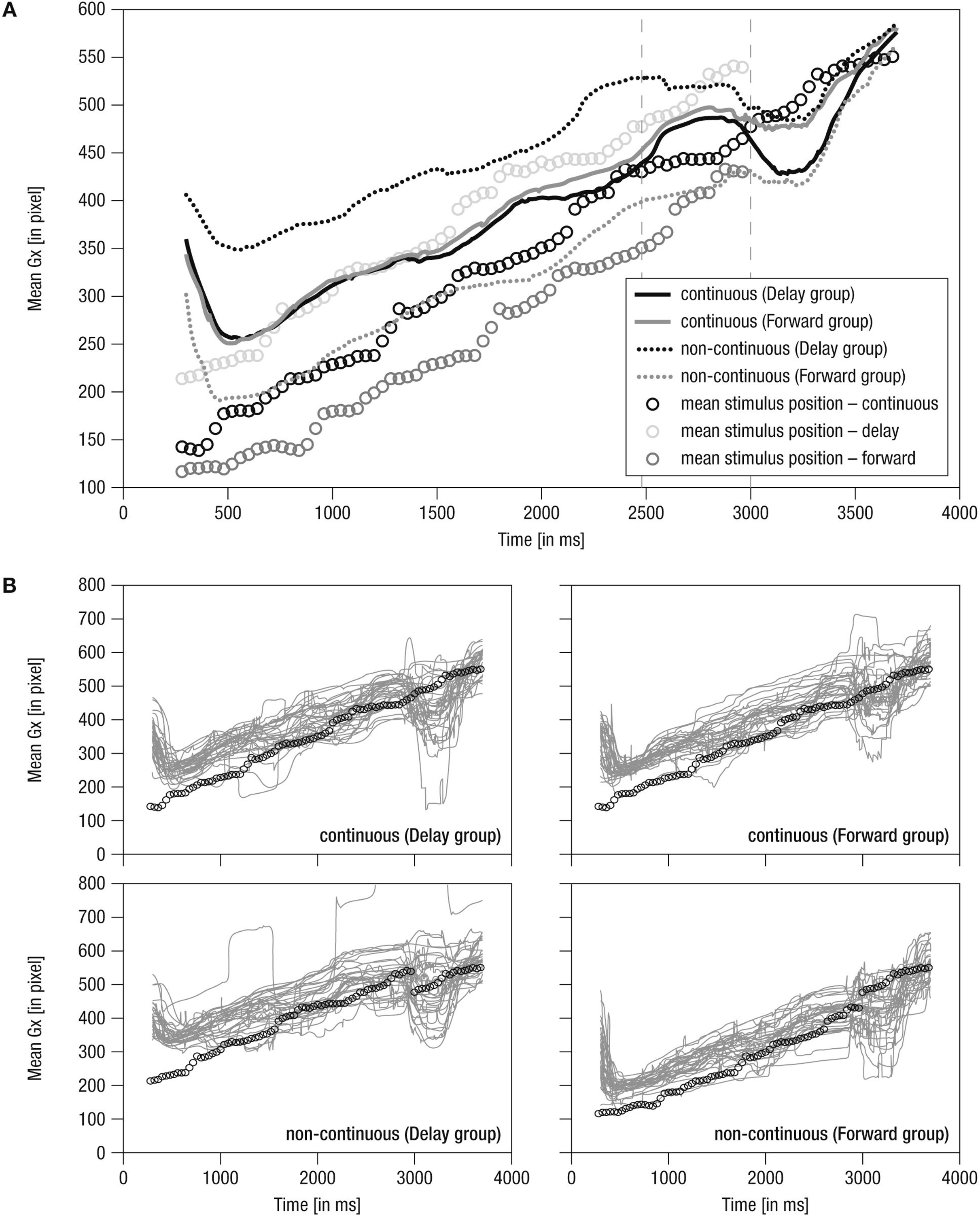
Mean horizontal gaze positions over time. (**A**) Grand averaged horizontal gaze positions over time. Lines: Solid – continuous, Dotted – non-continuous movement, Black – Delay group, Gray – Forward group, Vertical dashed – occlusion on-and offset. (**B**) Single averaged horizontal gaze positions over time (gray). Note that circles indicate mean stimulus position over time for the respective condition. Prior to occlusion, circles are horizontally shifted by ± 500 ms due to stimulus design. Gaze positions in continuous conditions closely match because the stimulus was identical. As the stimulus was not visible during occlusion (i.e., 2480–3000 ms), here, circles indicate imaginary continuation of the movement. Following occlusion (i.e., 3000–4000 ms), only circles for the continuous condition are plotted as the stimulus was identical in all conditions.

For statistical analysis, within subjects, the *mean distance* as well as the *variance in distance* between gaze and stimulus position were calculated for each trial across predefined 500 ms time windows for each phase of the trial (i.e., the last 500 ms of the pre-occlusion, the 500 ms of the occlusion, and the first 500 ms of the post-occlusion phase), and resulting means and variances, respectively, were averaged per condition. The two measures reveal different aspects of viewing behavior: Mean distance represents the average gaze position relative to the target position, and was thus taken to reflect tracking *accuracy*. Variance in distance represents the average fluctuation in tracking behavior, and was thus taken to reflect tracking *consistency* (i.e., whether tracking was rather consistent or random across infants).

#### 2.3.2 EEG data

##### 2.3.2.1 Recording and pre-processing

EEG was recorded continuously with a BrainAmp DC amplifier (BrainProducts GmbH, Gilching, Germany) from 32 active electrodes (actiCap by BrainProducts) inserted into a soft elastic cap according to the 10-20-system (EASYCAP GmbH, Herrsching, Germany). During recording, the right mastoid electrode served as reference and the left mastoid was recorded as an additional channel. Ground was placed at location AFz. Impedances were kept below 20 kΩ during preparation. The EEG was recorded with an analog pass-band of 0.1 to 250 Hz and digitized with a sampling rate of 1000 Hz.

Prior to EEG-preprocessing, based on behavioral coding of video-recordings, trials were discarded if infants *(a)* did not attend to the total duration of stimulus presentation and *(b)* produced limb movement that could be seen as part of imitative crawling. The latter criterion was chosen because we were interested in brain activity related to action observation but not to imitation. Furthermore, using Vision Analyzer 2 (Brain Products) for visual inspection, EEG trials were discarded which comprised broken channels or extreme/untypical artifacts (i.e., extensive movements). To this end, remaining EEG data were segmented into 4700 ms long epochs (from -2700 ms to 2000 ms relative to the onset of occlusion). Subsequent preprocessing and analyses were conducted using the FieldTrip (developed at the F.C. Donders Centre for Cognitive Neuroimaging, Nijmegen, The Netherlands; http://www2.ru.nl/fcdonders/fieldtrip/, Oostenveld et al., 2011) and custom-made routines operated in MATLAB 7.10.0 (MathWorks Inc., Natick, MA, USA).

Data were cleared of stereotypic artifacts using Independent Component Analysis (ICA; Jung et al., 2000). Specifically, ICs representing eye blinks, saccades, muscle activity, or instrumental noise were visually identified and discarded from further analysis by a trained rater (CB). To this end, all selected segments across all conditions were concatenated within subjects, filtered (high pass 1 Hz, low pass 100 Hz, 6^th^-order Butterworth-filter), and subjected to an extended infomax ICA (Bell and Sejnowski, 1995). A DFT-filter as implemented in FieldTrip was used to suppress line-noise. Decisions for rejection were based on integrated information from the ICs topography, power spectrum, event-related potentials (ERPs) as well as individual trials and the distribution of the IC over trials. Rejected ICs were in accordance with previous reports on typical artifacts in EEG data when stimulus presentation elicited eye-movements in a passive viewing paradigm (e.g., Plöchl et al., 2012).

All subsequent analyses were carried out in sensor space, based on the back-projection of the non-artifact ICs. Previously identified broken channels were interpolated after ICA-cleaning. Cleaned data was re-referenced to the mathematically linked mastoids, filtered (high pass 1 Hz, low pass 30 Hz, 6^th^-order Butterworth-filter), and segmented into 4000 ms epochs according to the onset of occlusion (-2480 ms to 1520 ms). For each single trial, the offset was removed by subtracting the average of the total epoch.

Rhythmic neural activity was analyzed by means of fast Fourier transformation (FFT) using an individualized data approach taking idiosyncrasies into account (Nesselroade et al., 2007). That is, we identified the individual peak frequency at the individual peak electrode in a given electrode cluster and frequency range (Doppelmayr et al., 1998;Werkle-Bergner et al., 2009). In line with the literature, *frontal theta* activity, considered as reflecting mnemonic processing (see Saby and Marshall, 2012 for a review), was defined as oscillatory activity within 4–6 Hz at frontal electrodes F3, Fz, F4, FC1, and FC2 (Orekhova et al., 1999;Orekhova et al., 2006). *Central alpha* activity, assumed to indicate sensorimotor simulation (for a review, see Marshall and Meltzoff, 2011), was defined as oscillatory activity within 6–9 Hz at central electrodes FC1, FC2, C3, Cz, C4, CP1, and CP2 (Stroganova et al., 1999;Marshall et al., 2002).

To detect individual peak frequencies, the spectral power distribution between 1 Hz and 20 Hz at each electrode was estimated by means of fast Fourier transformation (FFT) across all trials and phases (i.e., from -2480 ms to 1520 ms with regard to occlusion onset). Each trial was zero-padded to 10 s and tapered with a Hanning window to achieve a frequency resolution of 0.1 Hz. The power spectra were corrected for the 1/f trend inherent in scalp EEG data to facilitate the detection of spectral peaks (Demanuele et al., 2007;He et al., 2010). When no IPF was detected, the missing values were interpolated with the mean of all detected peaks to preserve comparable samples for the EEG measures. There was one missing value for frontal theta and central alpha each. These missings were not detected in the same participants across EEG measures.

##### 2.3.2.2 FFT analysis

For analyses of modulations in rhythmic neural activity, *FFT* was performed separately for each phase of the trial (i.e., pre-occlusion, occlusion, post-occlusion). As the phases (i.e., pre-occlusion, occlusion, post-occlusion) of each trial varied in length, the data were again zero-padded to 10 sec prior to FFT calculation, resulting in a common frequency resolution of 0.1 Hz. Power values for each phase of the trial and experimental condition were extracted for each participant at the respective individual preak frequency and electrode after averaging across trials within participants. For each condition, data were collapsed across movement directions (i.e., left to right and right to left) to obtain enough trials for statistical comparison. As the distribution of power values was skewed, data were log-transformed prior to the analysis^1^.

#### 2.4 Statistical analysis and qualitative description

To provide rich information on infants’ tracking behavior over the course of the stimulus movement, mean horizontal gaze positions as well as mean horizontal distance in gaze and stimulus positions over time were described qualitatively. In addition, statistical analyses were done using SPSS 15.0 (SPSS Inc., 1989–2006, USA). Specifically, mixed effects repeated-measures ANOVAs with a between-subject factor *Group* (Delay group vs. Forward group) and the within-subjects factors *Phase* (pre-occlusion vs. occlusion vs. post-occlusion phase) and *Time* (continuous vs. non-continuous) were carried out separately for each measure of eye movement (i.e., mean distance, variance in distance) and rhythmic neural activity (i.e., frontal theta, central alpha). Including Phase makes it possible to check that differences in dependent variables occur only after the time-course manipulation was introduced, namely during the post-occlusion phase. Partial eta squared, *η*_*p*_^*^2^*^, is reported as an estimate of the effect size. Greenhouse-Geisser corrections were applied if the assumption of sphericity was violated. As group sizes were equal, ANOVA was assumed to be robust towards violation of the assumption of homogeneity. Significant effects were followed up by separate Bonferroni-corrected ANOVAs or *t*-tests.

## 3 Results

### 3.1 Eye-tracking data

#### 3.1.1 Qualitative description of gaze positions over time

Mean horizontal gaze positions over time are shown in Figure 3.

*(1)* During the *pre-occlusion* phase, a decrease in horizontal gaze positions until 500 ms after trial onset indicates a slow orientation reaction. When infants were finally ‘on’ the stimulus, movement was tracked comparably across experimental groups and conditions in close relation to the stimulus position (Figure 3A). Note that in the forwarded/delayed conditions the stimulus depicted a movement that started500 ms earlier/later in the movement sequence than in the continuous conditions, and the crawling infant was thus at slightly different positions across conditions throughout the pre-occlusion phase (see Figure 2). Accordingly, gaze positions were about 150 pixels further backward in forwarded (see gray dotted line in Figure 3A) and further forwarded in delayed conditions (see black dotted line in Figure 3A) compared to continuous conditions.

*(2)* During the *occlusion* phase, general tracking behavior continued in accordance with the stimulus trajectory presented during the pre-occlusion phase. Towards the occlusion offset, the difference between non-continuous conditions reduced about 50 pixels, possibly indicating adaptation to non-matching stimulus reappearance in repeated/block stimulus presentation.

*(3)* At the *post-occlusion* onset, distinct tracking patterns emerged: In the case of continuous movement in the Delay group, infants’ gaze positions were reduced for about 50 pixels; that is, infants gazed opposite the movement direction (solid black line in Figure 3A). This was followed by catching-up with the stimulus movement (i.e., steep increase in horizontal gaze positions). All conditions were tracked comparably towards the end of the trial (i.e., at 3500 ms at about pixel 550). Note that visual input was identical in all conditions during the post-occlusion phase but did not match the continued time course of the pre-occlusion input in non-continuous continuations (i.e., delayed/forwarded). Hence, infants quickly caught up with the stimulus in response to manipulated continuations.

Notably, the grand averages reflected the individual data (Figure 3B) suggesting that tracking was rather consistent across infants. In sum, average raw gaze positions over time indicate that infants were sensitive to manipulations in the timing of observed movements.

#### 3.1.2 Qualitative description of distance in gaze and stimulus position over time

The average horizontal distance in gaze and stimulus position over time is shown in Figure 4.

**Figure 4.**
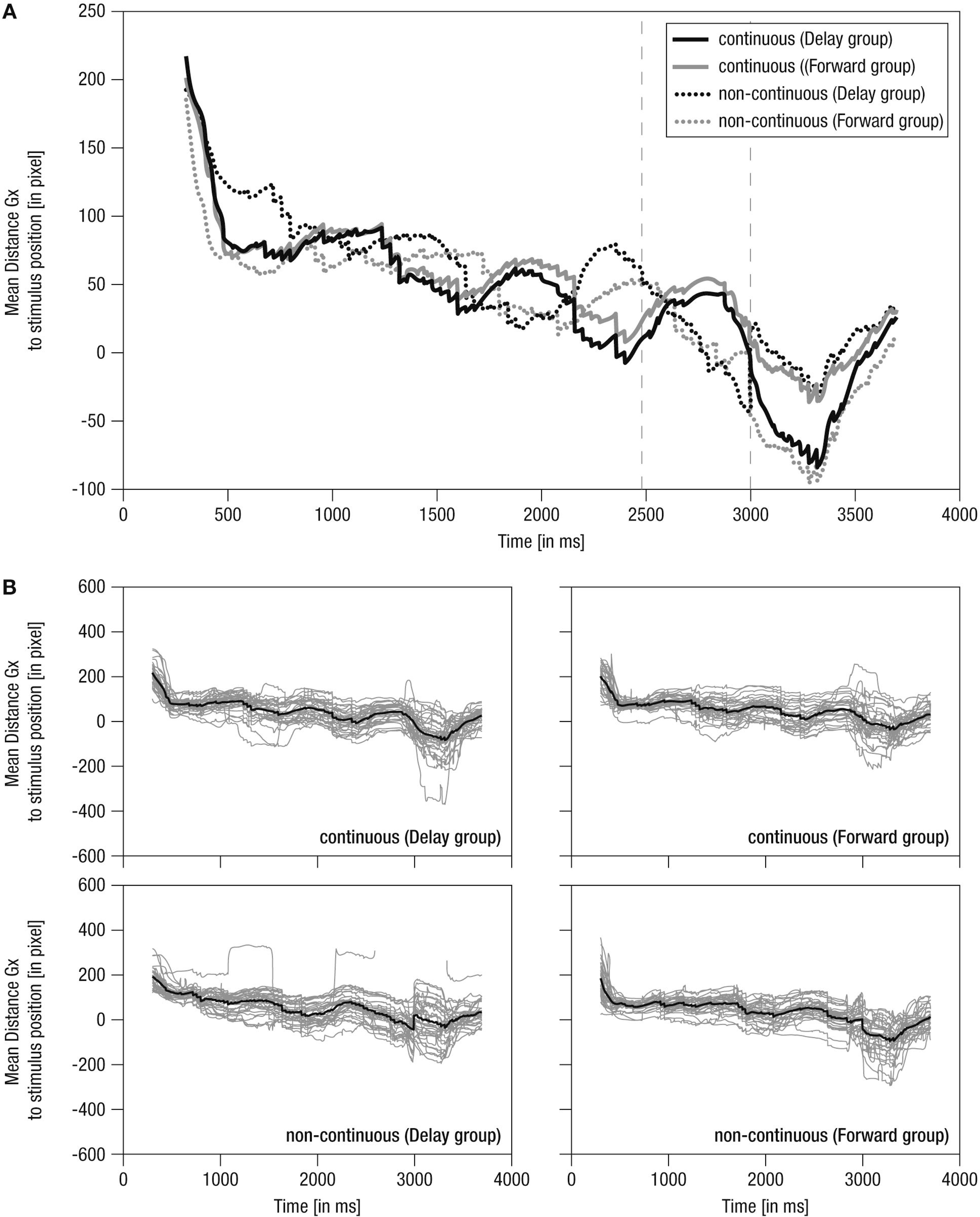
Mean horizontal distance between gaze positions and mean stimulus positions over time. (**A**) Grand averaged distance. Gx – raw gaze points on x-dimension. Lines: Solid – continuous, Dotted – non-continuous, Black – Delay group, Gray – Forward group; Vertical dashed – occlusion on-and offset. (**B**) Single averaged distance (gray) including respective grand average (black). Note the average stimulus dimensions of 279 pixel.

*(1)* During the *pre-occlusion* phase, both continuous and non-continuous movements were tracked in accordance with the non-linear dynamics of the crawling movement (Figure 4A). Specifically, positive scores indicate that infants preferably tracked the front to middle parts of the baby stimulus with decreasing scores (i.e., about 50 pixels over 2000 ms) when approaching the occlusion phase. This may indicate adaptation to the transient full-screen occlusion of the stimulus movement always occurring 2480 ms post stimulus-onset.

*(2)* During the *occlusion* phase, in continuous conditions, the cyclic tracking pattern was continued, indicating that infants stayed on the stimulus although it was hidden. In contrast, in non-continuous conditions, distance scores distinctively decreased about 100 pixels (i.e., looking opposing the hidden target’s implied movement direction) in *delayed* movement (i.e., converging to the reappearance position) and slightly decreased about 50 pixels in *forwarded* movement (i.e., diverging from the reappearance position). Nevertheless, infants were still ‘on’ the target in non-continuous conditions, yet on mid to rear parts of it. Hence, though movement manipulation could be detected following occlusion, infants apparently expected a certain continuation during occlusion, possibly due to repeated/blocked presentation of conditions.

*(3)* At the *post-occlusion* onset, tracking of continuous and non-continuous continuations differed between the experimental groups: In the *Delay group*, continuous movement resulted in a pronounced decrease in distance scores (i.e., about 100 pixels, thus looking opposite the movement direction) until the gaze was positioned on rear parts of the stimulus, whereas delayed movement resulted in a small decrease (i.e., about 40 pixels) until the gaze was positioned at the mean stimulus position. In contrast, in the *Forward group*, continuous movement resulted in only a small decrease (i.e., about 40 pixels) towards the mean stimulus position, whereas forwarded movement resulted in a pronounced decrease (i.e., about 100 pixels) towards rear parts of the stimulus. Hence, continuous movement was apparently not always perceived as time-matching continuation. Finally, following a steep increase in distance scores, all conditions were tracked comparably at about 50 pixels mean distance (i.e., at front parts of stimulus) 700 ms post occlusion-offset, showing that infants quickly caught up with the actual stimulus movement.

Like mean horizontal gaze positions, grand averages of mean horizontal distance in gaze and stimulus positions were representative of individual data, which were actually highly systematic across conditions and individuals (Figure 4B) highlighting that tracking behavior was rather consistent across participants. Overall, these results indicate that infants were able to detect slight temporal shifts in the continuation of transiently occluded movements.

#### 3.1.3 Statistical analysis of mean distance per phase

To analyze the *mean distance* as a marker for tracking accuracy in 500 ms time windows before, during, and following occlusion, a mixed effects repeated-measures ANOVA was performed. The results showed a significant main effect of the within-subjects factor *(a)* Phase (*F*_(1.6, 97.9)_ = 130.25, *p* = .000, *η*_*p*_^*^2^*^ = .68). Furthermore, there were significant interaction effects for *(b)* Phase and Time (*F*_(1.6, 97.1)_ = 4.59, *p* = .012, *η*_*p*_^*^2^*^ = .07), *(c)* Time and Group (*F*_(1, 61)_ = 10.37, *p* = .002, *η*_*p*_^*^2^*^ = .15), and *(d)* Phase, Time, and Group (*F*_(1.6, 97.1)_ = 17.1, *p* = .000, *η*_*p*_^*^2^*^ = .22). No further effects were observed (*F* < 3.06, *p* > .085). Figure 5 provides an overview of the results for mean distance and variance in distance.

**Figure 5.**
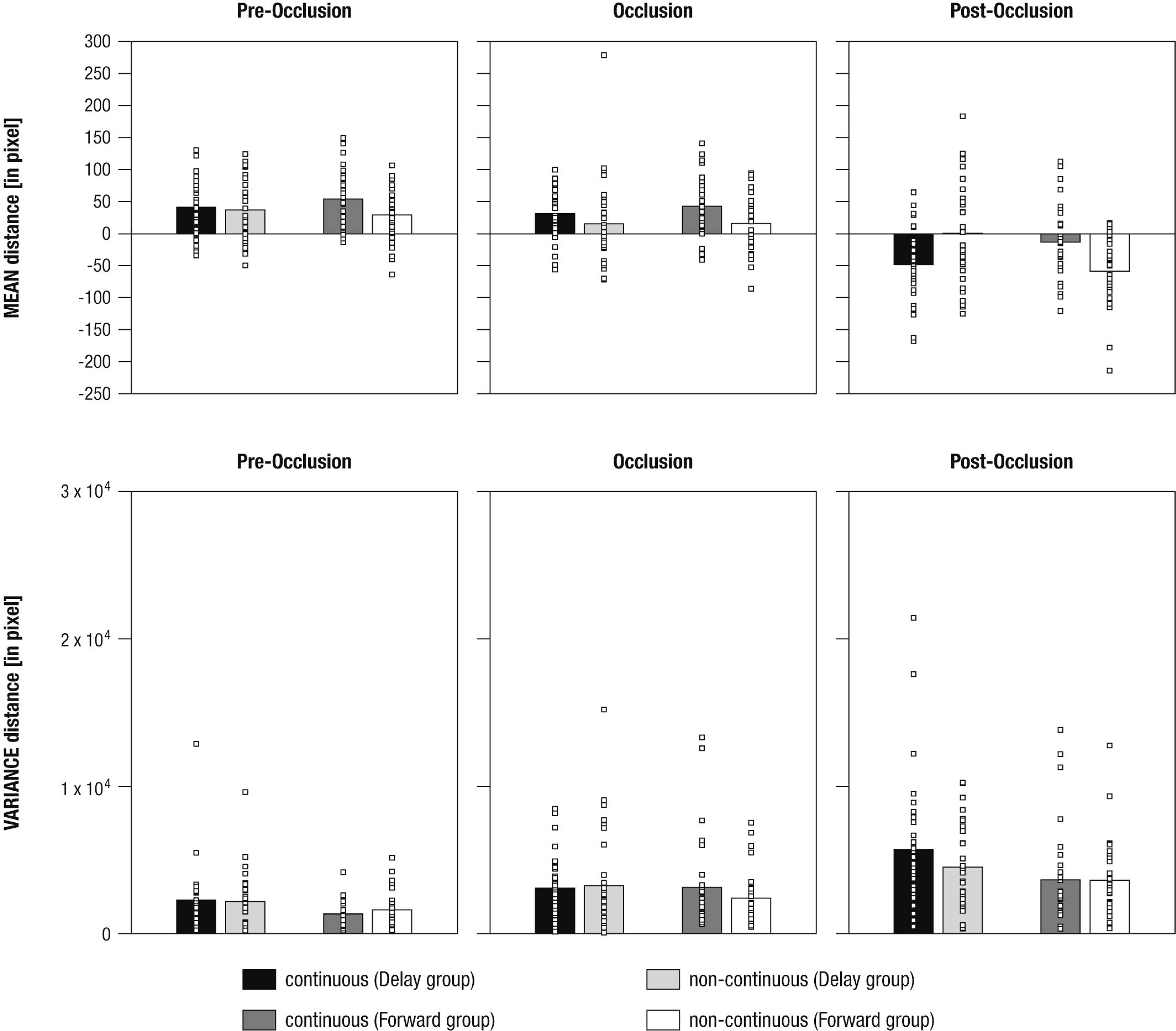
Mean differences in mean distance (upper panel) and variance in distance (lower panel) between gaze positions and mean stimulus positions shown separately for experimental conditions (i.e., continuous in the Delay group, continuous in the Forward group, non-continuous in the Delay group, non-continuous in the Forward group), and phases (i.e., pre-occlusion, occlusion, and post-occlusion). Squares indicate single cases to demonstrate the distribution within the sample.

To evaluate the (*d*) three-way interaction effect, a total of six paired-sample *t*-tests were performed, separately per levels of Group and Phase. The results showed that, during *post-occlusion*, the *Delay group* tracked continuous movements (*M* = -47.88, *SE* = 9.98) at more rear parts than non-continuous movements (*M* = 0.9, *SE* = 13.8; *t*_(31)_ *=* -3.25, *p* = .003; pre-occlusion: *t*_(31)_ *=*. 54, *p =*. 595; occlusion: *t*_(31)_ *=* 1.51, *p =*. 142), whereas the *Forward group* tracked continuous movements (*M* = -12.91, *SE* = 10.34) more frontally than non-continuous movements (*M* = -58.18, *SE* = 9.92; *t*_(31)_ *=* 3.69, *p* = .001; pre-occlusion: *t*_(30)_ *=* 2.1, *p =*. 03; occlusion: *t*_(30)_ *=* 2.0, *p =*. 05).

In sum, these results indicate that infants differentiated continuous from non-continuous movements following occlusion. However, as already indicated in the qualitative description of average distance over time (see *3.1.2*), continuous movement was apparently not tracked similarly across experimental groups: Corresponding to our hypotheses infants in the Forward group tracked continuous movements more accurately but undershot forwarded continuations. Counter to expectations, infants in the Delay group did not overshoot delayed, but undershot continuous movements.

#### 3.1.4 Statistical analysis of variance in distance per phase

To analyze the *variance in distance* as a marker of tracking consistency in 500 ms time windows before, during, and following occlusion, a mixed-effects repeated-measures ANOVA was calculated. This revealed significant main effects of the within-subjects factor *(a)* Phase (*F*_(1.7, 104.2)_ = 24.72, *p* = .000, *η*_*p*_^*^2^*^ = .29) and the between-subjects factor *(b)* Group (*F*_(1, 61)_ = 4.69, *p* = .034, *η*_*p*_^*^2^*^ = .07). No further effects were found (all *F* < 2.25, all *p* > .110).

Using paired-sample t-tests to follow up on the main effect of *(a)* Phase indicated that variance in distance was highest during post-occlusion (*M* = 4369.1, *SE* = 369.07; all *t*_(62)_ > 4.31, all *p* = .000). Variance in distance was also higher during occlusion (*M* = 2958.11, *SE* = 295.13) compared to pre-occlusion (*M* = 1842.46, *SE* = 189.36, *t*_(62)_ *= 3*.54, *p* = .001).

To follow-up on the main effect of *(b)* Group, an unpaired t-test showed that variance in distance was higher in the Delay group (*M* = 3487.29, *SE* = 314.94) than in the Forward group (*M* = 2611.92, *SE* = 250.87; *t*_(61)_ > 2.16, *p* = .034).

In sum, variance in distance increased due to transient occlusions. In addition, tracking was less consistent overall when infants watched continuous and delayed crawling versus continuous and forwarded crawling.

Taken together, both qualitative and statistical analyses of gazing behavior combine to provide a consistent picture: Results indicate that infants detected slight manipulations of the time course of an observed movement. Specifically, infants watching continuous and forwarded movements produced a tracking pattern consistent with the hypothesis of internal real-time simulation of observed movements during a transient occlusion (Graf et al., 2007). In contrast, infants watching continuous and delayed movements, albeit discriminating both conditions, produced a tracking pattern suggesting that real-time representations were not always precise (enough) or possibly altered by further processing (e.g., learned expectations across repeated presentations).

### 3.2 EEG data

#### 3.2.1 Frontal theta activity

To analyze mnemonic contributions to time-course representations, a mixed effects repeated-measures ANOVA was calculated for frontal theta activity. Results showed a significant main effect of Phase (*F*_(1.55, 2.06)_ = 5.72, *p* = .009, *η*_*p*_^*^2^*^ = .57) without evidence for further effects (all *F* < 1.41; all *p >*. 250). Figure 6 provides an overview of the EEG results. Hence, counter to expectations, no differential activation of frontal theta activity was found, indicating that the manipulation of the time course of ongoing movement did not elicit differential demands on memory processes.

**Figure 6.**
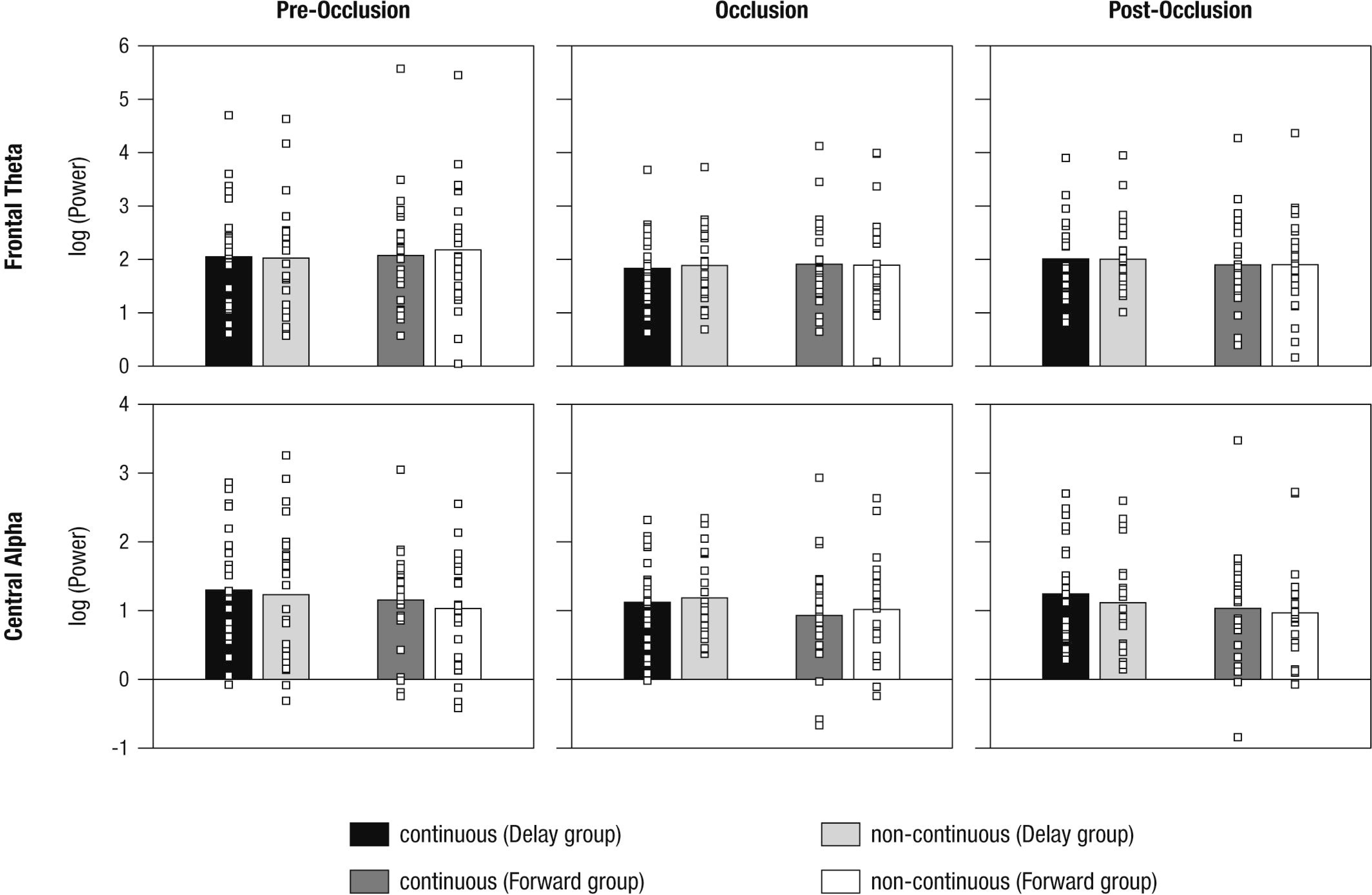
Mean power differences between experimental conditions (i.e., continuous in the Delay group, continuous in the Forward group, non-continuous in the Delay group, non-continuous in the Forward group) and phases (i.e., pre-occlusion, occlusion, and post-occlusion) for frontal theta and central alpha activity. Squares indicate single cases to demonstrate the distribution within the sample.

#### 3.2.2 Central alpha activity

To analyze contributions from sensorimotor simulation to time-course representations, a mixed effects repeated-measures ANOVA was performed for central alpha activity. A significant interaction effect of Phase and Time occurred (*F*_(1.9, 91.5)_ = 3.61, *p* = .031, *η*_*p*_^*^2^*^ = .07). No further effects were observed (all *F* < 2.14, all *p >*. 123).

As also implied by the small effect size, follow-up repeated measures ANOVAs separately per level of Phase, did not yield significant effects (all *F* < 2.64, all *p >*. 110). From the inspection of results as displayed in Figure 6 it may be concluded that, during post-occlusion, central alpha activity was lower for *non-continuous* than for continuous movements. Hence, in line with our hypothesis, our findings suggest that the cortical sensorimotor system is involved when infants render real-time simulations of transiently occluded movements that are within their motor repertoire.

## 4 Discussion

This study explored the internal representation of the *time course* of observed movement. To this end, 10-months-old crawling infants watched videos of a same-aged crawling baby that was transiently occluded and reappeared in a time-matching (i.e., continuous) or non-matching (i.e., delayed vs. forwarded) manner. To tap mnemonic and sensorimotor contributions to time-course representations, eye movement and rhythmic neural activity were simultaneously measured. First, the results suggest that sensorimotor functions were recruited more during the perception of non-matching continuations following occlusion. In contrast, there was no evidence for a differential role of mnemonic functions for time-course representations. Secondly, eye movements differentiated between time-matching and non-matching continuations following occlusion indicating a high sensitivity to the movements’ time course. In sum, we conclude that 10-month-old infants generate internal movement representations that reflect the timing of observed movements. This corresponds to the internal real-time simulation account of action observation (Graf et al., 2007).

### 4.1 Eye movements are sensitive to the time course of movements

To investigate infants’ sensitivity to the time course of observed movements, we assessed eye-tracking patterns in response to a transiently occluded human movement. Our findings showed that 10-month-old infants distinguished between temporally matching and temporally shifted (i.e., delayed vs. forwarded) continuations following occlusion as demonstrated by differences in the mean distance in gaze and stimulus position.

Previous studies have indicated that 4- to 7-month-old infants are largely insensitive to a manipulation in the timing of an object’s motion during occlusion, in that temporal violations were only detected in extreme cases (i.e., instantaneous reappearance on the other side of an occluding board; Wilcox and Schweinle, 2003;Bremner et al., 2005). Only at the age of 2 years did toddlers’ searching behavior demonstrate an understanding for the relation between time, velocity, and distance when a train went through a tunnel (Möhring et al., 2012). Adults were more accurate in identifying one of multiple moving objects when the objects instantaneously disappeared and reappeared at the position they had vanished or even before that position but not when the objects reappeared at a linearly extrapolated position along their movement trajectory (Keane and Pylyshyn, 2006). Nevertheless, the present study illustrates 10-month-old crawling infants’ sensitivity to slight temporal shifts when observing videos of a crawling baby.

We can think of at least three possible reasons why infants in the present study were able to detect temporal changes. First, manipulation in the timing of an object’s motion, as carried out in previous infant studies (Wilcox and Schweinle, 2003;Bremner et al., 2005), might be processed differently than manipulation in the timing of a *human action* because body form and dynamics offer rich information on, for instance, changes in velocity or direction (Hernik et al., 2014;Wronski and Daum, 2014). This notion corresponds to studies in adults showing that occluded human actions are internally simulated in real-time (Graf et al., 2007;Parkinson et al., 2012;Springer et al., 2013). Moreover, actions with natural human kinematics have been found to be more accurately predicted than those with artificial ones (Stadler et al., 2012). Similarly, proficient motor experience has been shown to enhance prediction of reappearance positions (Stapel et al., 2016).

Second, previous studies predominantly investigated object motion during the first months of life only (e.g., von Hofsten et al., 1998;Wilcox and Schweinle, 2003;Bremner et al., 2005), whereas the present study investigated human motion in 10-month-olds. Though the *developmental trajectory* of time-course representation is poorly understood to date, one may assume that older infants are better at solving temporal shifts in movement, irrespective of the observed target.

Third, in most studies, data on infants’ gazing behavior are reduced to a selection of putatively relevant aspects, for example, to overall looking time following habituation (e.g., Bremner et al., 2005) or to predictive looking at the end of an observed action (e.g., Henrichs et al., 2014). While the data reduction approach has doubtlessly provided interesting information, it may also have prevented researchers from discovering further early capabilities (see also Roberts, 2004). Here, *rich data* on the gaze progression over time were analyzed, demonstrating 10-month-old infants’ spatiotemporal sensitivity while observing continuous and time-manipulated human movement that was within their own motor repertoire.

### 4.2 Sensorimotor processing is sensitive to the time course of movements

To explore the neural basis of internal real-time processing, we assessed rhythmic neural oscillations related to mnemonic (i.e., frontal theta) and sensorimotor processing (i.e., central alpha) while infants were observing movements that were either time-matching or non-matching following a transient occlusion.

*Frontal theta* activity did not differ between time-matching and non-matching continuations. Thus, we found no evidence that slight time-course manipulations in ongoing movement pose differential mnemonic demands on 10-month-old infants. Frontal theta, as measured here, is thought to implement a neural accumulator (Bland and Oddie, 2001;van Vugt et al., 2012) assisting in maintaining and integrating extracted information across time and space (e.g., Miller and Cohen, 2001;Simons and Spiers, 2003). Correspondingly, it has been shown that, in 10-month-old infants, mnemonic functions support the binding of pre-and post-occlusion movement input into a coherent and unified percept (Bache et al., 2015). The present finding however modifies the notion of mnemonic contributions, suggesting that precise temporal representations for movement integration may not be provided by mnemonic functions alone (Wilson, 2001;Coppe et al., 2010).

For *central alpha* activity, we found a significant interaction effect between the timing of movement (i.e., continuous vs. non-continuous) across the phases of the trial (i.e., pre-occlusion, occlusion, post-occlusion). Although it was not possible to discern the direction of the effect in follow-up analyses, inspection of Figure 6 suggests differences between time-matching and non-matching continuations following occlusion. Central alpha, as observed here, has been associated with sensorimotor simulation during movement observation (Cochin et al., 1999;Muthukumaraswamy et al., 2004;Marshall et al., 2011). Therefore, the present findings indicate sensorimotor involvement in the internal simulation of the timing of human movement. This interpretation is also supported by concurrent findings on eye movements (as described above), suggesting that the non-reliable differences in neural activity may not be due to infants’ lacking capabilities to detect differences in movements’ time courses.

Behavioral and neuroimaging studies in adults and infants suggest a crucial role of sensorimotor brain areas in timed internal simulation (e.g., Schubotz and von Cramon, 2002;Graf et al., 2007;Southgate et al., 2009;Stadler et al., 2011;Cross et al., 2012;Elsner et al., 2013;Springer et al., 2013;Stapel et al., 2016). Such a predictive function of the motor system may allow reduction of the processing delay in sensory–motor loops, which pose a fundamental challenge to proactive control of perception and behavior (e.g., Blakemore and Frith, 2005;Schubotz, 2007). However, simulating sensorimotor consequences in real-time may not (yet) be fast, stable, or precise enough in 10-month-old crawlers observing a crawling movement (see Wolpert and Flanagan, 2001).

### 4.3 Further considerations

Effects of either delayed or forwarded continuations were most obvious when comparing time-matching continuations between the two groups (Delayed and Forwarded). We assumed that, if occluded movement was internally simulated in real-time, infants would undershoot reappearance positions in forwarded continuations and undershoot them in delayed continuations, whereas infants would accurately track reappearance positons in continuous movements. Results showed that infants alternately watching continuous and forwarded movements produced a tracking pattern consistent with this hypothesis. However, infants alternately watching continuous and delayed movements undershot time-matching continuations and overshot delayed continuations. In fact, the tracking patterns of both experimental groups were found to be unexpectedly overlapping (see Figure 4): Infants watching continuous and delayed movements tracked the *continuous* movement in a similar way as infants watching continuous and forwarded movements tracked the *forwarded* movement. Vice versa, infants watching continuous and forwarded movements pursued the *continuous* movement in a similar way as infants watching continuous and delayed movements pursued the *delayed* movement. Moreover, tracking was less consistent across infants, when infants watched continuous and delayed continuations in contrast to continuous and forwarded continuations. Note however, that the variation between conditions is a between subject comparison, i.e., two different groups of subjects performed delayed and forwarded conditions.

Though illustrating infants’ remarkable sensitivity to an action’s time course, these findings cannot solely be explained in terms of internal real-time processing. We can, however, only speculate as to which processes may have contributed to the pattern of results.

First, the present findings suggest that delayed and forwarded time-shifts in observed human action are not processed similarly (Bremner et al., 2005;Striano et al., 2006). This corresponds to adult studies showing that adults judged the continuation of actions following an occlusion to be continuous when it was in fact slightly delayed while slightly forwarded continuations were judged correctly as forwarded (e.g., Sparenberg et al., 2012). Switching from tracking external motion to internally representing motion may be costly and may thus lead to misaligned internal processing (Sparenberg et al., 2012; see also Mitrani and Dimitrov, 1978). In line with this notion, it is not obvious whether infants in the present study detected delayed continuations as manipulated in time. Future studies are needed to pinpoint the threshold at which time-matching and non-matching continuations are experienced as equal to determine potential *switching costs* early in life.

Second, the present findings may indicate that continuous movements are not always perceived as such (see also Adler et al., 2008). An influence of the stimulus context on action perception may be explained in accordance with *priming* effects (e.g., Pavlova and Sokolov, 2000). For example, when adults first performed a seemingly unrelated motor task (e.g., arm movement) and later observed movements corresponding to the motor task (i.e., arm movement) and non-corresponding (i.e., leg movement), the evaluation of the timing of movement continuations following occlusion was facilitated in corresponding conditions (Springer et al., 2013). Priming during action observation has also been reported in infant populations (e.g., Daum and Gredeback, 2011). From this perspective, non-matching conditions here may have served as the prime altering the processing of the time-matching condition. Future studies may disentangle whether and how time-shifted movements can change the perception of alternately presented continuous movements.

Third, it is possible that expectations based on *learning* across the repeated/blocked presentation of conditions may have contributed to the present results. This may be assumed because infants seem to have adapted their gaze position according to the expected reappearance position when approaching the occlusion offset (see Figure 4). Specifically, they looked slightly further back in delayed and slightly further forward in forwarded movements. In addition, following occlusion, there was a tendency to undershoot movements irrespective of the actual condition, which may be interpreted as an overall conservative strategy to stay on the target following a transient full-screen occlusions (cf. Stapel et al., 2016). At the same time, differences in tracking following occlusion suggest that infants did not learn that the stimulus’ reappearance position was kept *identical* in all conditions (see *2.2*). Future studies should clarify whether and how learning may contribute to internal time-course representations when infants observe repetitive human movements.

There was a considerable drop-out on the level of trials and participants in both eye and brain measures. *High attrition rates* of 25–75% are commonly observed in EEG studies with mobile infant populations (see de Haan, 2007;for a meta-analysis see Stets et al., 2012). In eye-tracking studies with infants, drop-out on the level of trials and participants has not been documented consistently. Concurrent preparation of both EEG and eye-tracking reduces potential testing time and challenges infants’ compliance (e.g., see number of infants who could not be properly tested in *2.1*). Furthermore, both methods are sensitive to gross body and head movements that may result in a critical loss of data. In addition, eye-tracking is sensitive to repeated, persistent, and substantial changes in the position of the eyes (due to changes of head and/or body position), and measurement quality decreases over time in head-free recording (Holmqvist et al., 2011). At the same time, multiple repetition of the stimulus material is required for EEG to reduce noise in the signal. Therefore, it seems reasonable to assume comparable drop-out rates for eye-tracking and EEG data, and, potentially, overall higher attrition in simultaneous measurement in comparison to single measurement of either brain or eye data. Furthermore, not all participants can be expected to contribute (enough) data to both measures.

As a *consequence of high attrition*, it was not possible here to directly relate EEG and eye-tracking measures (see also Stapel et al., 2010). Furthermore, it cannot be excluded that attrition was selective for infants who complied better with testing requirements (e.g., Marshall et al., 2009) restricting the generalizability of effects. Moreover, due to infrequent and random contribution of data (see Figure 1), a systematic analysis of tracking over time (i.e., within and across blocks) was not conducted, because it would have required reducing the number of available trials and participants substantially.

From a methodological perspective, eye movements elicited during action perception add a source of artifacts to the EEG measurement potentially distorting the results. In adults, it has been shown that eye tracking data measured simultaneously with EEG can be used to identify and correct for those artifacts (e.g., Dimigen et al., 2011;Plöchl et al., 2012). In contrast, in infants, automated approaches to clean EEG of stereotypic artifacts are lacking. Here, we visually identified ICs representing eye movement related artifacts. Even though the ICA produced meaningful results in accordance with the adult literature, we cannot be certain whether artifacts were sufficiently removed in all data because eye and brain data could not directly be related as discussed above.

### 4.4 Conclusion

In this study, an experimental paradigm previously used to investigate internal real-time processing during action perception in adults (e.g., Graf et al., 2007) was successfully adapted and applied to an infant population. We found that 10-month-old crawlers are able to detect slight manipulations of the timing of observed crawling movements as reflected in infants’ tracking and neural patterns. This suggests a remarkable sensitivity to spatiotemporal information about external events early in life.

## Conflict of Interest Statement

The authors declare that the research was conducted in the absence of any commercial or financial relationships that could be construed as a potential conflict of interest.

1 Comparable results were obtained in non-log-transformed data after exclusion of outliers (> mean ±3*SD).

## Author Contributions

CB, AS, WS, FK, and UL conceived and designed the study, CB collected the data, CB, HN, and MWB analyzed and interpreted the data, CB drafted the manuscript, all authors revised the work and approved the final version for publication.

## Acknowledgements

This research was supported by the Max Planck Research Network for the Cognitive and Neurosciences (Maxnet *Cognition*) and funded by the Max Planck Society. The study was conducted in partial fulfillment of the doctoral dissertation of CB. CB received training and financial support from the International Max Planck Research School on the Life Course (LIFE, http://www.imprs-life.mpg.de). We cordially thank the infants and their parents for participating in this study and our student assistants for their support in data collection, coding and preprocessing. We further wish to thank Berndt Wischnewski for technical assistance and Julia Delius for editorial help.

## References

Adamson, L.B., and Frick, J.E. (2003). The still face: a history of a shared experimental paradigm. Infancy 4, 451–473. doi:10.1207/S15327078IN0404_01

Adler, S.A., Haith, M.M., Arehart, D.M., and Lanthier, E.C. (2008). Infants' visual expectations and the processing of time. J. Cogn.Dev. 9, 1’25. doi:10.1080/15248370701836568

Bache, C., Kopp, F., Springer, A., Stadler, W., Lindenberger, U., and Werkle-Bergner, M. (2015). Rhythmic neural activity indicates the contribution of attention and memory to the processing of occluded movements in 10-month-old infants. Int. J. Psychophysiol. 98, 201’212. doi:10.1016/j.ijpsycho.2015.09.003

Baddeley, A.D., and Hitch, G.J. (1974). “Working memory,” in The Psychology of Motivation and Learning, ed. G.H. Bower. (New York: Academic Press), 47-89.

Bell, A.J., and Sejnowski, T.J. (1995). An information-maximization approach to blind separation and blind deconvolution. Neural Comput. 7, 1129’1159. doi:10.1162/neco.1995.7.6.1129

Bennett, S.J., and Barnes, G.R. (2003). Human ocular pursuit during the transient disappearance of a visual target. J. Neurophysiol. 90, 2504’2520. doi:10.1152/jn.01145.2002

Blakemore, S.J., and Frith, C. (2005). The role of motor contagion in the prediction of action. Neuropsychologia 43, 260’267. doi:10.1016/j.neuropsychologia.2004.11.012

Bland, B.H., and Oddie, S.D. (2001). Theta band oscillation and synchrony in the hippocampal formation and associated structures: the case for its role in sensorimotor integration. Behav. Brain Res. 127, 119’136. doi:10.1016/S0166-4328(01)00358-8

Bosco, G., Delle Monache, S., and Lacquaniti, F. (2012). Catching what we can't see: manual interception of occluded fly-ball trajectories. PLoS One 7, e49381. doi:10.1371/journal.pone.0049381

Bremner, J.G., Johnson, S.P., Slater, A., Mason, U., Foster, K., Cheshire, A., and Spring, J. (2005). Conditions for young infants' perception of object trajectories. Child Dev. 76, 1029’1043. doi:10.1111/j.1467-8624.2005.00895.x

Canfield, R.L., and Haith, M.M. (1991). Young infants visual expectations for symmetrical and asymmetric stimulus sequences. Dev. Psych. 27, 198’208. doi:10.1037/0012-1649.27.2.198

Cochin, S., Barthelemy, C., Roux, S., and Martineau, J. (1999). Observation and execution of movement: similarities demonstrated by quantified electroencephalography. Eur. J. Neurosci. 11, 1839’1842. doi:10.1046/j.1460-9568.1999.00598.x

Coppe, S., De Xivry, J.J., Missal, M., and Lefevre, P. (2010). Biological motion influences the visuomotor transformation for smooth pursuit eye movements. Vision Res. 50, 2721’2728. doi:10.1016/j.visres.2010.08.009

Cross, E.S., Liepelt, R., Hamilton, A.F., Parkinson, J., Ramsey, R., Stadler, W., and Prinz, W.(2012). Robotic movement preferentially engages the action observation network. Hum. Brain Mapp. 33, 2238’2254. doi:10.1002/hbm.21361

Cuevas, K., Cannon, E.N., Yoo, K., and Fox, N.A. (2014). The Infant EEG Mu Rhythm:Methodological Considerations and Best Practices. Developmental Review 34, 26’43. doi:10.1016/j.dr.2013.12.001

Daum, M.M., and Gredeback, G. (2011). The development of grasping comprehension in infancy: covert shifts of attention caused by referential actions. Exp. Brain Res. 208, 297’307. doi:10.1007/s00221-010-2479-9

De Haan, M. (2007). “Visual attention and recognition memory in infancy,” in Infant EEG and Event-Related Potentials, ed. M. De Haan. (New York: Psychology Press), 101’138.

Demanuele, C., James, C.J., and Sonuga-Barke, E.J. (2007). Distinguishing low frequency oscillations within the 1/f spectral behaviour of electromagnetic brain signals. Beh. Brain Funct. 3, 62. doi:1744-9081-3-62 [pii] 10.1186/1744-9081-3-62

Dimigen, O., Sommer, W., Hohlfeld, A., Jacobs, A.M., and Kliegl, R. (2011). Coregistration of eye movements and EEG in natural reading: analyses and review. J Exp Psychol Gen 140, 552’572. doi:10.1037/a0023885

Doppelmayr, M., Klimesch, W., Pachinger, T., and Ripper, B. (1998). Individual differences in brain dynamics: important implications for the calculation of event-related band power. Biol. Cybern. 79, 49’57. doi:10.1007/s004220050457

Elsner, C., D'ausilio, A., Gredebäck, G., Falck-Ytter, T., and Fadiga, L. (2013). The motor cortex is causally related to predictive eye movements during action observation. Neuropsychologia 51, 488’492. doi:10.1016/j.neuropsychologia.2012.12.007

Falck-Ytter, T., Gredebäck, G., and Von Hofsten, C. (2006). Infants predict other people's action goals. Nat. Neurosci. 9, 878’879. doi:10.1038/nn1729

Flanagan, J.R., and Johansson, R.S. (2003). Action plans used in action observation. Nature 242, 769’771. doi:10.1038/nature01861

Graf, M., Reitzner, B., Corves, C., Casile, A., Giese, M., and Prinz, W. (2007). Predicting point-light actions in real-time. NeuroImage 36, 22’32. doi:10.1016/j.neuroimage.2007.03.017

Gredebäck, G., and Melinder, A. (2010). Infants' understanding of everyday social interactions: a dual process account. Cognition 114, 197’206. doi:10.1016/j.cognition.2009.09.004

Green, D., Kochukhova, O., and Gredebäck, G. (2014). Extrapolation and direct matching mediate anticipation in infancy. Infant Beh. Dev. 37, 111’118. doi:10.1016/j.infbeh.2013.12.002

Haith, M.M. (2004). Progress and standardization in eye movement work with human infants.Infancy 6, 257’265. doi:10.1207/s15327078in0602_6

He, B.J., Zempel, J.M., Snyder, A.Z., and Raichle, M.E. (2010). The temporal structures and functional significance of scale-free brain activity. Neuron 66, 353’369. doi:S0896-6273(10)00291-6 [pii]10.1016/j.neuron.2010.04.020

Hebb, D.O. (1949/2009). The organization of behavior - a neuropsychological theory. New York:Wiley.

Henrichs, I., Elsner, C., Elsner, B., Wilkinson, N., and Gredebäck, G. (2014). Goal certainty modulates infants' goal-directed gaze shifts. Dev. Psych. 50. doi:10.1037/a0032664

Hernik, M., Fearon, P., and Csibra, G. (2014). Action anticipation in human infants reveals assumptions about anteroposterior body-structure and action. Proc. R. Soc. Lond. [Biol] 281, 20133205. doi:10.1098/rspb.2013.3205

Holmqvist, K., Nyström, M., Andersson, R., Dewhurst, R., Jarodzka, H., and Van De Weijer, J.(2011). Eye Tracking: A Comprehensive Guide to Methods and Measures. Oxford: Oxford University Press.

Hommel, B., Musseler, J., Aschersleben, G., and Prinz, W. (2001). The theory of event codin (TEC): a framework for perception and action planning. Behav. Brain Sci. 24, 849’937. doi:10.1017/S0140525X01000103

Hunnius, S., and Bekkering, H. (2010). The early development of object knowledge: a study of infants' visual anticipations during action observation. Dev. Psych. 46, 446’454. doi:10.1037/a0016543

Jacobs, J., and Kahana, M.J. (2010). Direct brain recordings fuel advances in cognitive electrophysiology. Trends Cogn. Sci. 14, 162’171. doi:10.1016/j.tics.2010.01.005

Jung, T.P., Makeig, S., Humphries, C., Lee, T.W., Mckeown, M.J., Iragui, V., and Sejnowski, T.J.(2000). Removing electroencephalographic artifacts by blind source separation. Psychophysiology 37, 163’178. doi:10.1111/1469-8986.3720163

Keane, B.P., and Pylyshyn, Z.W. (2006). Is motion extrapolation employed in multiple object tracking? Tracking as a low-level, non-predictive function. Cogn. Psychol. 52, 346’368. doi:10.1016/j.cogpsych.2005.12.001

Kilner, J.M., Friston, K.J., and Frith, C.D. (2007). Predictive coding: an account of the mirror neuron system. Cogn. Process. 8, 159’166. doi:10.1007/s10339-007-0170-2

Komogortsev, O.V., Gobert, D.V., Jayarathna, S., Koh, D.H., and Gowda, S.M. (2010). Standardization of automated analyses of oculomotor fixation and saccadic behaviors. IEEE Trans. Biomed. Eng. 57, 2635’2645. doi:10.1109/Tbme.2010.2057429

Kwon, M.K., Luck, S.J., and Oakes, L.M. (2014). Visual short-term memory for complex objects in 6-and 8-month-old infants. Child Dev. 85, 564’577. doi:10.1111/Cdev.12161

Lisman, J.E., and Jensen, O. (2013). The theta-gamma neural code. Neuron 77, 1002’1016. doi:10.1016/j.neuron.2013.03.007

Marshall, P.J., Bar-Haim, Y., and Fox, N.A. (2002). Development of the EEG from 5 months to 4 years of age. Clin. Neurophysiol. 113, 1199’1208. doi:10.1016/S1388-2457(02)00163-3

Marshall, P.J., and Meltzoff, A.N. (2011). Neural mirroring systems: exploring the EEG mu rhythm in human infancy. Dev. Cogn. Neurosci. 1, 110’123. doi:10.1016/j.dcn.2010.09.001

Marshall, P.J., Reeb, B.C., and Fox, N.A. (2009). Electrophysiological responses to auditory novelty in temperamentally different 9-month-old infants. Dev. Sci. 12, 568’582. doi:10.1111/j.1467-7687.2008.00808.x

Marshall, P.J., Young, T., and Meltzoff, A.N. (2011). Neural correlates of action observation and execution in 14-month-old infants: an event-related EEG desynchronization study. Dev. Sci. 14, 474’480. doi:10.1111/j.1467-7687.2010.00991.x

Miller, E.K., and Cohen, J.D. (2001). An integrative theory of prefrontal cortex function. Annu. Rev. Neurosci. 24, 167’202. doi:10.1146/annurev.neuro.24.1.167

Mitrani, L., and Dimitrov, G. (1978). Pursuit eye movements of a disappearing moving target Vision Res. 18, 537’539. doi:10.1016/0042-6989(78)90199-2

Möhring, W., Cacchione, T., and Bertin, E. (2012). On the origin of the understanding of time, speed, and distance interrelations. Infant Beh. Dev. 35, 22’28. doi:10.1016/j.infbeh.2011.09.008

Muthukumaraswamy, S.D., Johnson, B.W., and Mcnair, N.A. (2004). Mu rhythm modulation during observation of an object-directed grasp. Cogn. Brain Res. 19, 195’201. doi:10.1016/j.cogbrainres.2003.12.001

Nesselroade, J.R., Gerstorf, D., Hardy, S.A., and Ram, N. (2007). Idiographic filters for psychological constructs. Measurement, 217–235. doi:10.1080/15366360701741807

Oostenveld, R., Fries, P., Maris, E., and Schoffelen, J.-M. (2011). FieldTrip: open source software for advanced analysis of MEG, EEG, and invasive electrophysiological data. Comput. Intell. Neurosci. 2011, 1. doi:10.1155/2011/156869

Orekhova, E.V., Stroganova, T.A., and Posikera, I.N. (1999). Theta synchronization during sustained anticipatory attention in infants over the second half of the first year of life. Int. J. Psychophysiol. 32, 151’172. doi:10.1016/S0167-8760(99)00011-2

Orekhova, E.V., Stroganova, T.A., Posikera, I.N., and Elam, M. (2006). EEG theta rhythm in infants and preschool children. Clin. Neurophysiol. 117, 1047’1062. doi:10.1016/j.clinph.2005.12.027

Parkinson, J., Springer, A., and Prinz, W. (2012). Before, during and after you disappear: aspects of timing and dynamic updating of the real-time action simulation of human motions. Psychol. Res. 76, 421’433. doi:10.1007/s00426-012-0422-3

Pavlova, M., and Sokolov, A. (2000). Orientation specificity in biological motion perception. Percept Psychophys 62, 889’899. doi:10.3758/Bf03212075

Pelphrey, K.A., and Reznick, J.S. (2002). Working memory in infancy. Adv. Child Dev. Behav. 31, 173’227. doi:10.1016/S0065-2407(03)31005-5

Plöchl, M., Ossandon, J.P., and König, P. (2012). Combining EEG and eye tracking: identification, characterization, and correction of eye movement artifacts in electroencephalographic data. Frontiers in human neuroscience 6, 1’23. doi:

Roberts, S. (2004). Self-experimentation as a source of new ideas: ten examples about sleep, mood, health, and weight. Behav. Brain Sci. 27, 227’262. doi:10.1017/S0140525X04000068

Rosander, K., and Von Hofsten, C. (2004). Infants' emerging ability to represent occluded object motion. Cognition 91, 1’22. doi:10.1016/S0010-0277(03)00166-5

Rosander, K., and Von Hofsten, C. (2011). Predictive gaze shifts elicited during observed and performed actions in 10-month-old infants and adults. Neuropsychologia 49, 2911’2917. doi:10.1016/j.neuropsychologia.2011.06.018

Saby, J.N., and Marshall, P.J. (2012). The utility of EEG band power analysis in the study of infancy and early childhood. Dev. Neuropsychol. 37, 253’273. doi:10.1080/87565641.2011.614663

Schneider, M., Heine, A., Thaler, V., Torbeyns, J., De Smedt, B., Verschaffel, L., Jakobs, A.M., and Stern, E. (2008). A validation of eyemovements as a measure of elementary school children’s developing number sense. Cognitive Dev. 23, 409’422. doi:10.1016/j.cogdev.2008.07.002

Schubotz, R.I. (2007). Prediction of external events with our motor system: towards a new framework. Trends Cogn. Sci. 11, 211’218. doi:10.1016/j.tics.2007.02.006

Schubotz, R.I., and Von Cramon, D.Y. (2002). Dynamic patterns make the premotor cortex interested in objects: influence of stimulus and task revealed by fMRI. Cogn. Brain Res. 14, 357’369. doi:10.1016/S0926-6410(02)00138-6

Simons, J.S., and Spiers, H.J. (2003). Prefrontal and medial temporal lobe interactions in long-term memory. Nat. Rev. Neurosci. 4, 637’648. doi:10.1038/nrn1178

Southgate, V., Johnson, M.H., Karoui, I.E., and Csibra, G. (2010). Motor system activation reveals infants' on-line prediction of others' goals. Psychol. Sci. 21, 355’359. doi:10.1177/0956797610362058

Southgate, V., Johnson, M.H., Osborne, T., and Csibra, G. (2009). Predictive motor activation during action observation in human infants. Biol. Lett. 5, 769’772. doi:10.1098/rsbl.2009.0474

Sparenberg, P., Springer, A., and Prinz, W. (2012). Predicting others' actions: evidence for a constant time delay in action simulation. Psychol. Res. 76, 41’49. doi:10.1007/s00426-011-0321-z

Springer, A., Brandstädter, S., and Prinz, W. (2013). Dynamic simulation and static matching for action prediction: evidence from body part priming. Cogn. Sci. 37, 936’952. doi:10.1111/cogs.12044

Stadler, W., Schubotz, R.I., Von Cramon, D.Y., Springer, A., Graf, M., and Prinz, W. (2011). Predicting and memorizing observed action: differential premotor cortex involvement. Hum. Brain Mapp. 32, 677’687. doi:10.1002/hbm.20949

Stadler, W., Springer, A., Parkinson, J., and Prinz, W. (2012). Movement kinematics affect action prediction: comparing human to non-human point-light actions. Psychol. Res. 76, 395’406. doi:10.1007/s00426-012-0431-2

Stampe, D.M. (1993). Heuristic filtering and reliable calibration methods for video-based pupil-tracking systems. Behav. Res. Methods Instrum. Comput. 25, 137’142. doi:10.3758/Bf03204486

Stapel, J.C., Hunnius, S., Meyer, M., and Bekkering, H. (2016). Motor system contribution to action prediction: temporal accuracy depends on motor experience. Cognition 148, 71’78. doi:10.1016/j.cognition.2015.12.007

Stapel, J.C., Hunnius, S., Van Elk, M., and Bekkering, H. (2010). Motor activation during observation of unusual versus ordinary actions in infancy. Soc. Neurosci. 5, 451’460. doi:10.1080/17470919.2010.490667

Stets, M., Stahl, D., and Reid, V.M. (2012). A meta-analysis investigating factors underlying attrition rates in infant ERP studies. Dev. Neuropsychol. 37, 226’252. doi:10.1080/87565641.2012.654867

Striano, T., Henning, A., and Stahl, D. (2006). Sensitivity to interpersonal timing at 3 and 6 months of age. Interaction Stud. 7, 251’271. doi:10.1075/is.7.2.08str

Stroganova, T.A., Orekhova, E.V., and Posikera, I.N. (1999). EEG alpha rhythm in infants. Clin. Neurophysiol. 110, 997’1012. doi:10.1016/S1388-2457(98)00009-1

Van Vugt, M.K., Simen, P., Nystrom, L.E., Holmes, P., and Cohen, J.D. (2012). EEG oscillations reveal neural correlates of evidence accumulation. Front. Neurosci. 6, 106. doi:10.3389/fnins.2012.00106

Von Hofsten, C., Vishton, P., Spelke, E.S., Feng, Q., and Rosander, K. (1998). Predictive action in infancy: tracking and reaching for moving objects. Cognition 67, 255’285. doi:10.1016/S0010-0277(98)00029-8

Wass, S.V., Forssman, L., and Leppanen, J.M. (2014). Robustness and precision: How data quality may influence key dependent variables in infant eye-tracker analyses. Infancy 19, 427’460. doi:10.1111/infa.12055

Wass, S.V., Smith, T.J., and Johnson, M.H. (2013). Parsing eye-tracking data of variable quality to provide accurate fixation duration estimates in infants and adults. Behav. Res. Methods 45, 229’250. doi:10.3758/s13428-012-0245-6

Werkle-Bergner, M., Shing, Y.L., Muller, V., Li, S.C., and Lindenberger, U. (2009). EEG gamma-band synchronization in visual coding from childhood to old age: evidence from evoked power and inter-trial phase locking. Clin. Neurophysiol. 120, 1291’1302. doi:10.1016/j.clinph.2009.04.012

Wilcox, T., and Schweinle, A. (2003). Infants' use of speed information to individuate objects in occlusion events. Infant Beh. Dev. 26, 253’282. doi:10.1016/S0163-6383(03)00021-3

Wilson, M. (2001). The case for sensorimotor coding in working memory. Psychon B Rev 8, 44’57. doi:10.3758/Bf03196138

Wolpert, D.M., and Flanagan, J.R. (2001). Motor prediction. Curr. Biol. 11, 729’732. doi:10.1016/S0960-9822(01)00432-8

Wronski, C., and Daum, M.M. (2014). Spatial orienting following dynamic cues in infancy: grasping hands versus inanimate objects. Dev. Psych. 50, 2020’2029. doi:10.1037/a0037155

